# Second Harmonic Generation Imaging Reveals Entanglement of Collagen Fibers in the Elephant Trunk Skin Dermis

**DOI:** 10.1101/2023.08.11.553031

**Authors:** Andrew K. Schulz, Magdalena Plotczyk, Sophia Sordilla, Madeline Boyle, Krishma Singal, Joy S. Reidenberg, David L. Hu, Claire A. Higgins

## Abstract

Form-function relationships often have tradeoffs: if a material is tough, it is often inflexible, and vice versa. This is particularly relevant for the elephant trunk, where the skin should be protective yet elastic. To investigate how this is achieved, we used classical histochemical staining and second harmonic generation microscopy to describe the morphology and composition of elephant trunk skin. We report structure at the macro and micro scales, from the thickness of the dermis to the interaction of 10 *μ*m thick collagen fibers. We analyzed several sites along the length of the trunk, to compare and contrast the dorsal-ventral and proximal-distal skin morphologies and compositions. We find the dorsal skin of the elephant trunk can have keratin armor layers over 2mm thick, which is nearly 100 times the thickness of the equivalent layer in human skin. We also found that the structural support layer (the dermis) of elephant trunk contains a distribution of collagen-I (COL1) fibers in both perpendicular and parallel arrangement. The bimodal distribution of collagen is seen across all portions of the trunk, and is dissimilar from that of human skin where one orientation dominates within a body site. We hypothesize that this distribution of COL1 in the elephant trunk allows both flexibility and load-bearing capabilities. Additionally, when viewing individual fiber interaction of 10 *μ*m thick collagen, we find the fiber crossings per unit volume are five times more common than in human skin, suggesting that the fibers are entangled. We surmise that these intriguing structures permit both flexibility and strength in the elephant trunk. The complex nature of the elephant skin may inspire the design of materials that can combine strength and flexibility.

## Introduction

Elephant trunks, octopus arms, and mammalian tongues are the three canonical examples of muscular hydrostats (Kier and Smith, 1985). The elephant trunk, the subject of this work, is extremely flexible and can extend by up to 25% in a telescopic manner allowing the elephant to reach distant objects (Schulz, Boyle, Boyle, Sordilla, Rincon, Hooper, Aubuchon, Reidenberg, Higgins, and Hu, 2022). The ventral side of the trunk contains oblique muscles that allows that part of the trunk to wrap around and grasp objects (Kier and Smith, 1985). It follows that the ventral surface of the trunk is often the primary point of contact between the trunk and the substrate during object manipulation (Dagenais, Hensman, Haechler, and Milinkovitch, 2021). The dorsal side of the trunk is not often utilized for grasping, and this surface of the trunk is more exposed to external mechanical forces and predators, potentially necessitating a more protective armor-like structure. To fulfill the different roles required of it, the skin on the elephant trunk is required to be flexible and tough at the same time.

Relatively little work has been conducted to observe and document the anatomy of elephant skin. In 1970, Spearman published a study discussing elephant skin’s basic anatomy, including insights about the different vibrissal hairs on the trunk (Spearman, 1970). More recently, biomechanical studies have made connections between the skin properties and an elephant’s ability to grasp and wrap its trunk around various objects, including barbells (Dagenais et al., 2021; Schulz, Reidenberg, Wu, Tang, Seleb, Mancebo, Elgart, and Hu, 2023). While the skin on the elephant body is cracked for thermoregulation (Martins, Bennett, Clavel, Groenewald, Hensman, Hoby, Joris, Manger, and Milinkovitch, 2018), the trunk, in contrast, has wrinkles and folds on its ventral and dorsal surfaces, respectively (Schulz et al., 2023). The structure also varies with position along the length of the trunk : the distal trunk skin (on both ventral and dorsal surfaces) is characterized by wrinkles, while the proximal dorsal trunk skin has folds. These differing skin characteristics enable the trunk to extend to reach faraway objects, with the dorsal surface stretching more than the ventral (Schulz et al., 2022).

In this work, we used both classical and newly developed microscopy techniques to investigate the structure of elephant trunk skin. We focused our analysis on collagen, a foundational protein that governs the structure of many body tissues, including muscle, blood vessels, and skin, and provides bio-inspiration across scales (Eder, Amini, and Fratzl, 2018). Collagen I (COL1) is the primary collagen found within the skin; it has a fibrillar structure and can therefore be detected with second harmonic generation (SHG) imaging. SHG is a nonlinear optical imaging technique that selectively detects noncentrosymmetric molecules, including type I and II collagen with no labelling (Boddupalli and Bratlie, 2015; Chen, Nadiarynkh, Plotnikov, and Campagnola, 2012).

SHG microscopy works by viewing the skin sample at a specific frequency that excites the fibrillar structure of COL1; the resulting image exhibits half the wavelength of the original wavelength used, hence the term “second harmonic.” The fibrillar structure of COL1 fibers allows the microscopy technique to detect COL1 in the tissue, while the resulting image is related to the amount of pre-strain on the COL1 fibers (Turcotte, Mattson, Wu, Zhang, and Lin, 2016). The SHG technique is label-free and therefore accrues less error compared to traditional histochemistry since there is not a chained sequence of staining that can vary based on the specific timing that segmented skin spends in various chemical baths(Haggerty, Wang, Dickinson, O’Malley, and Martin, 2014). The SHG technique is specific to collagen and does not pick up the other fiber structures, such as elastin or keratin that are present within the skin (Chen et al., 2012). In skin, COL1 networks are characterized by variations in fiber orientation, thickness, density, strain, and weaving with neighboring fibers -this last feature is a phenomenon known as entanglementDay, Zamani-Dahaj, Bozdag, Burnetti, Bingham, Conlin, Ratcliff, and Yunker (2023). Analysis of SHG images of skin allows quantification of all these variations in COL1 fibers.

We here used SHG to analyze COL1 architecture in elephant trunk skin. We conducted morphological and compositional analyses on skin samples along the trunk at several locations, including seven sites for SHG microscopy and eight for histochemical staining (**Figure 1**). We show key differences in collagen architecture along the length of the trunk, and differences between COL1 architecture in elephant and human skin.

**Figure 1:**
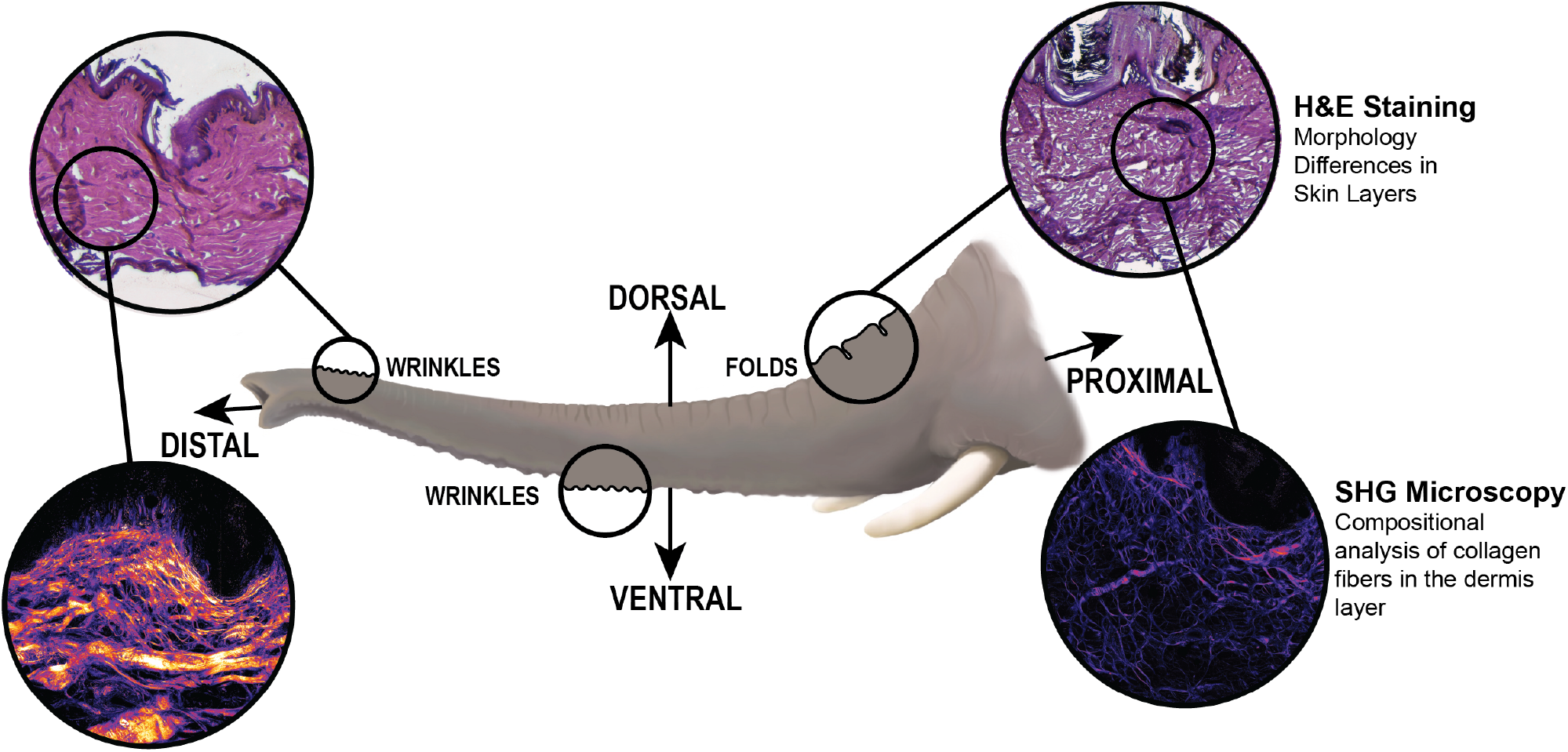
Schematic of the elephant trunk with experimental outputs from H&E Staining and SHG microscopy shown as insets.

## Experimental Methods

### Dissection of elephant trunk skin

Icahn School of Medicine at Mount Sinai, New York, provided access to a dissected frozen trunk from a 38-year-old female African elephant (*Loxodonta africana*) that initially lived in a Virginia zoo. The elephant was euthanized for health issues in 2011.

We accessed the trunk when it was on loan from the National Museum of Natural History (NMNH), Smithsonian Institution. The elephant’s body weight before death was approximately 4000 kg. The trunk was cut into several parts and initially stored in a freezer at −20°*C* until it was dissected in July 2016.

In March 2019, eight samples of the trunk skin were further dissected at the Icahn School of Medicine at Mount Sinai. These samples included five dorsal and three ventral samples ranging from the proximal to the distal end of the trunk. These samples were shipped on dry ice to Imperial College London by the Smithsonian Institute Collections Department as a scientific exchange between the two CITES-registered institutions. The Animal Plant and Health Agency in the UK (authorization number ITIMP19.0822) approved the tissue shipment. The samples were stored at Imperial College London at *−*80°C until embedding, sectioning, and imaging were conducted from January to March 2020.

### Histology and Morphometrics

The eight samples were further dissected to enable analysis in the trunk’s longitudinal direction. Samples were embedded in OCT (optimum cutting temperature) medium and 20 *μ*m-thick sections were cut on a cryostat (**Figure S1**). The tissue sections were stained using hematoxylin and eosin (H&E) and then imaged on a Zeiss inverted microscope at 3x magnification. Images were automatically segmented using the wand tool in FIJI (ImageJ) based on the stained color differences from H&E.

To quantify the thicknesses of each layer (the stratum corneum (SC), the viable epidermis (VE), and dermis (D) shown on **Figure 2**), a MATLAB script was used to divide each H&E image (1000 pixels wide) into vertical strips of one-pixel width. The pixels corresponding to each layer were counted and recorded. To compare samples, we reported the thickness for each layer, defined as the thickness of the layer divided by the sum of all layers (Table 1, **Figure 3**A, **Figure S2, Figure S3**).

**Table 1:** Table displaying the thickness of each skin layer in mm displayed in mean *±* standard deviation. Results of each layer are displayed as SC (**Figure 3**A), VE (**Figure S2**), and D (**Figure S3**).

| Dorsal Thickness |  |  |  |
| --- | --- | --- | --- |
| Distance from Tip | SC (mm) | VE (mm) | D (mm) |
| 3 | $0.39 \pm 0.11$ | $0.05 \pm 0.06$ | $0.8 \pm 0.097$ |
| 27 | $0.17 \pm 0.16$ | $0.36 \pm 0.21$ | $1.46 \pm 0.27$ |
| 81 | $0.39 \pm 0.27$ | $0.27 \pm 0.21$ | $6.6 \pm 0.28$ |
| 100 | $0.87 \pm 0.61$ | $0.58 \pm 0.51$ | $5.8 \pm 0.90$ |
| 133 | $1.83 \pm 0.68$ | $0.42 \pm 0.36$ | $5.44 \pm .64$ |
| Ventral Thickness |  |  |  |
| Distance from Tip | SC (mm) | VE (mm) | D (mm) |
| 27 | $0.29 \pm 0.32$ | $0.50 \pm 0.31$ | $2.4 \pm 0.44$ |
| 51 | $0.12 \pm 0.18$ | $0.32 \pm 0.49$ | $4.90 \pm 0.54$ |
| 133 | $0.34 \pm 0.22$ | $0.22 \pm 0.22$ | $4.19 \pm 0.23$ |

**Figure 2:**
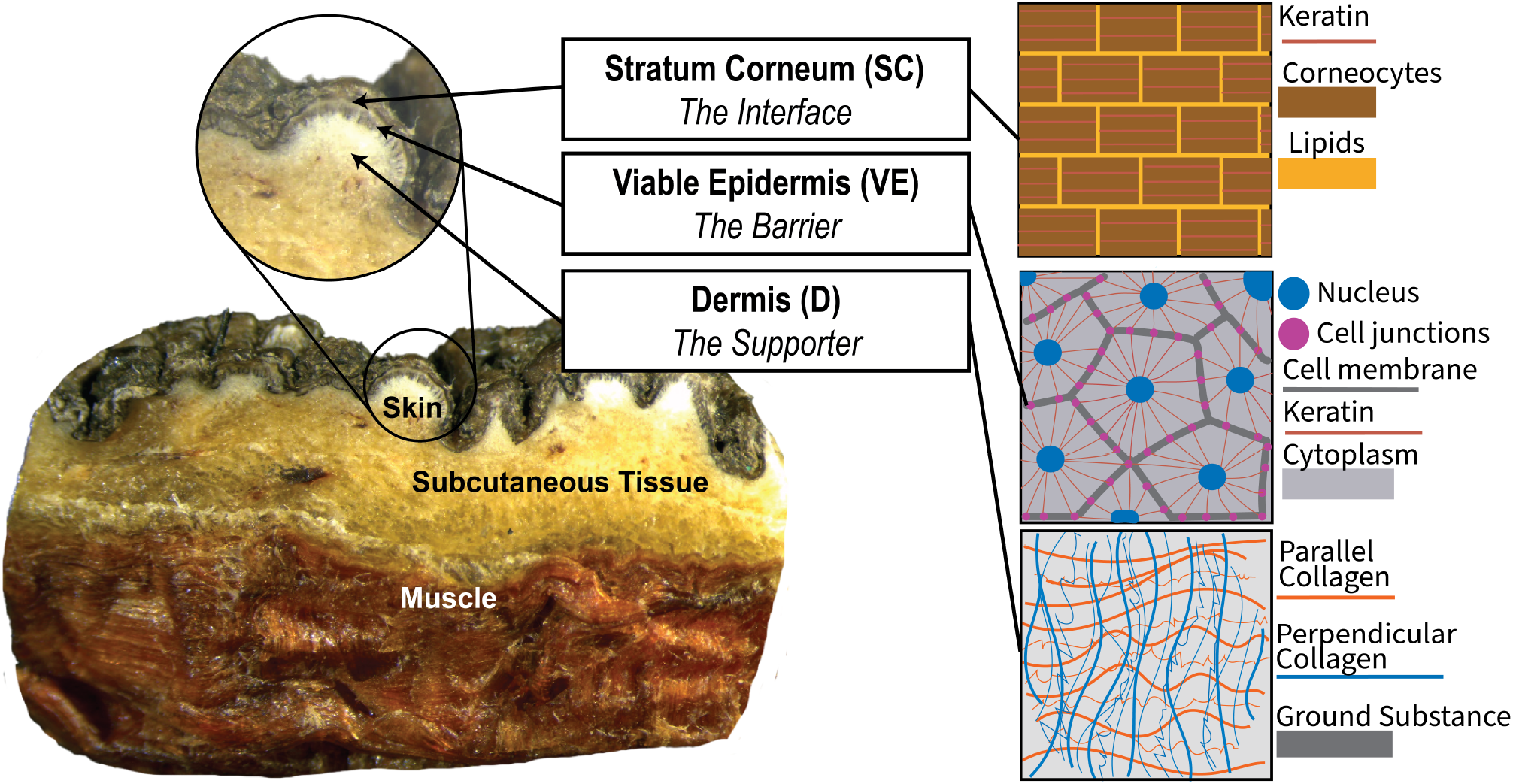
Macroscopic image of a cross section of elephant skin showing subcutaneous tissue and muscle. The skin layers are shown in a schematic of the Stratum Corneum (SC), Viable Epidermis (VE), and Dermis (D).

**Figure 3:**
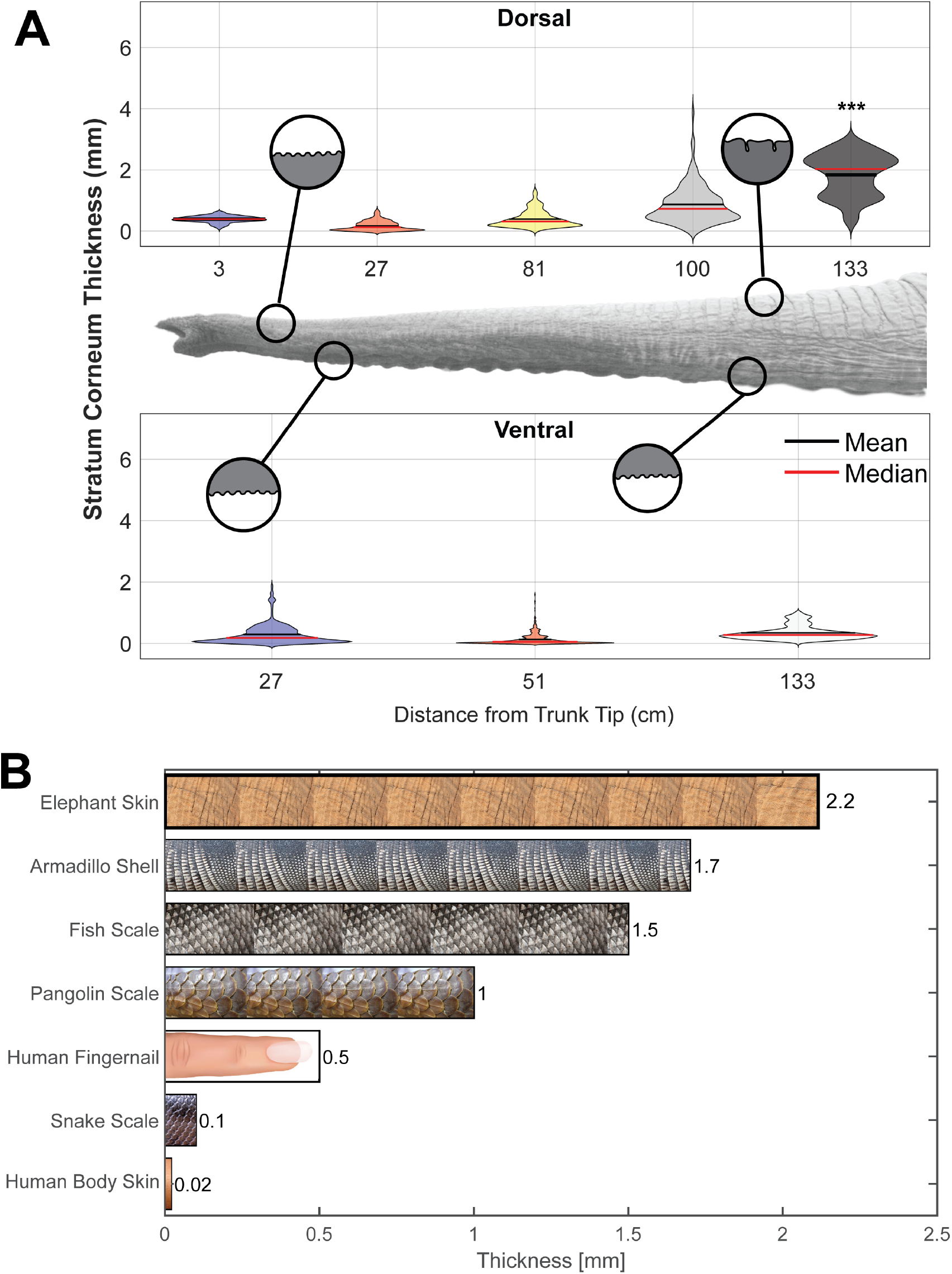
A) Relationship between Stratum corneum (SC) thickness and position on the trunk. The position is the distance from the trunk tip in cm. Stars indicate the statistical significance of the difference between dorsal and ventral sites: (^***^*p <* 0.001) B) Thickness of different dermal armors across species. Non-elephant data taken from (Bordoloi, 2021; Chintapalli et al., 2014; Han and Young, 2018; Wang et al., 2016; Wollina et al., 2001; Yang et al., 2019). Silhouettes and animal images taken from Adobe CC Images.

### Second Harmonic Generation

Samples embedded in OCT were sectioned at 100*μm* thickness for second-harmonic generation (SHG) imaging. Images were taken from an upright confocal microscope (Leica SP5) coupled to a Ti: Sapphire laser (Newport Spectra-Physics). Raw images were received as a stacked TIF file with 10 *μ*m between each image of the TIF file at a maximum of 255 nm with green luminescence. Stacks were then processed using a workflow in Fiji (ImageJ), including setting the minimum-maximum range to (0,4000), applying the blur filter (*σ*= 0.5), and subtracting background (rolling ball radius, 40 pixels). A machine-learning and segmenting open-source software, ilastik, was used to analyze the difference between fibers and background. Completed images are shown on **Figure 4**A-B.

**Figure 4:**
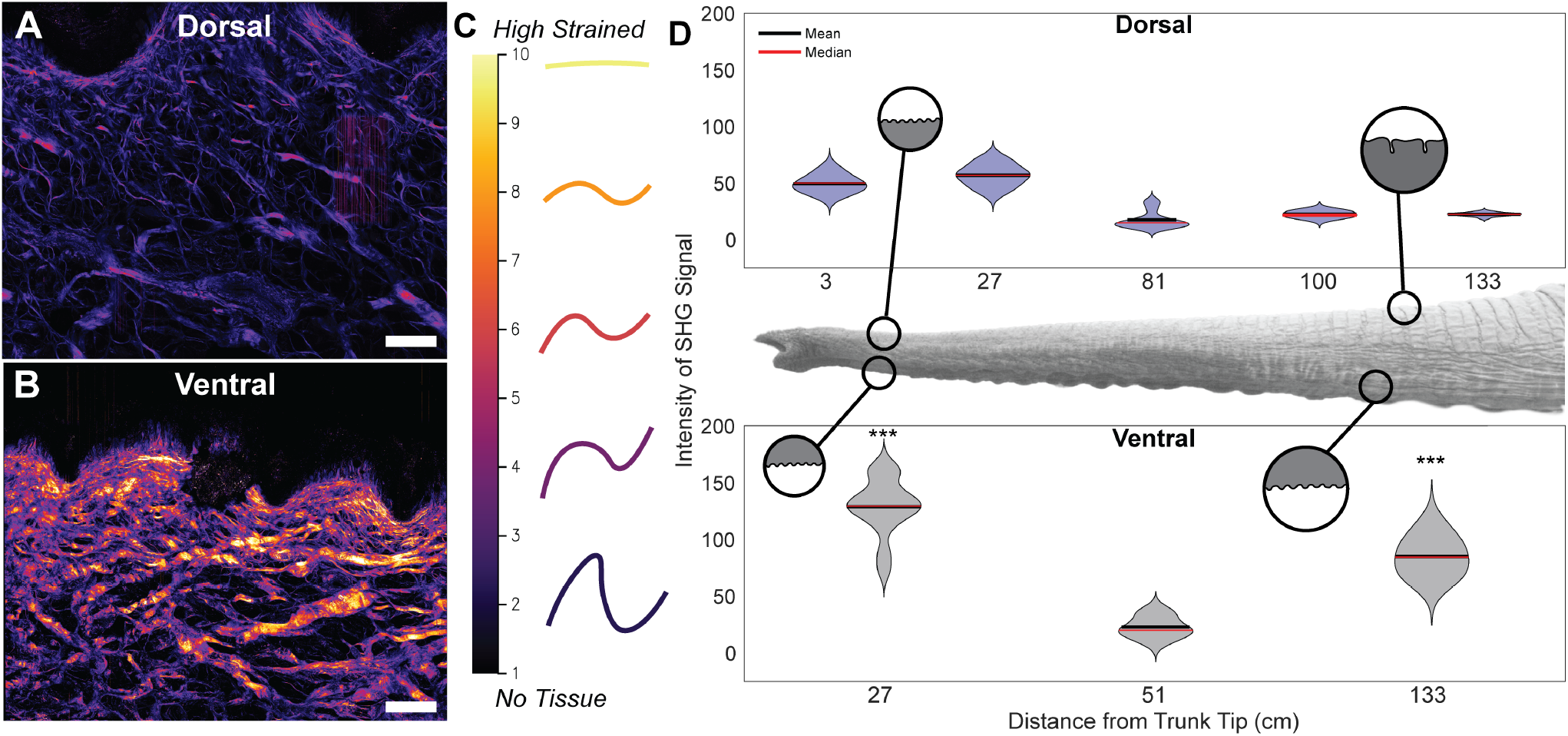
A-B) SHG stacked image of dorsal and ventral sections of the proximal trunk. C) Schematic displaying the relationship between the intensity of SHG in fibers and the indicative strain of a fiber. D) Relationship between SHG intensity and position on the trunk. Stars indicate the statistical significance of the difference between dorsal and ventral sites: (^***^ *p <* 0.001). Scale bars A,B: 100 *μ*m.

### Collagen Fiber Orientation and crossings

We used the open-source software CurveAlign to quantify the collagen fiber orientation in SHG images (Bredfeldt, Liu, Pehlke, Conklin, Szulczewski, Inman, Keely, Nowak, Mackie, and Eliceiri, 2014). Images were broken into regions of interest of size 600 *μ*m × 450 *μ*m with at least a 150 pixels distance from the boundary (**Figure 5**A). For this study, we only examined individual fibers instead of the entire fiber network.

**Figure 5:**
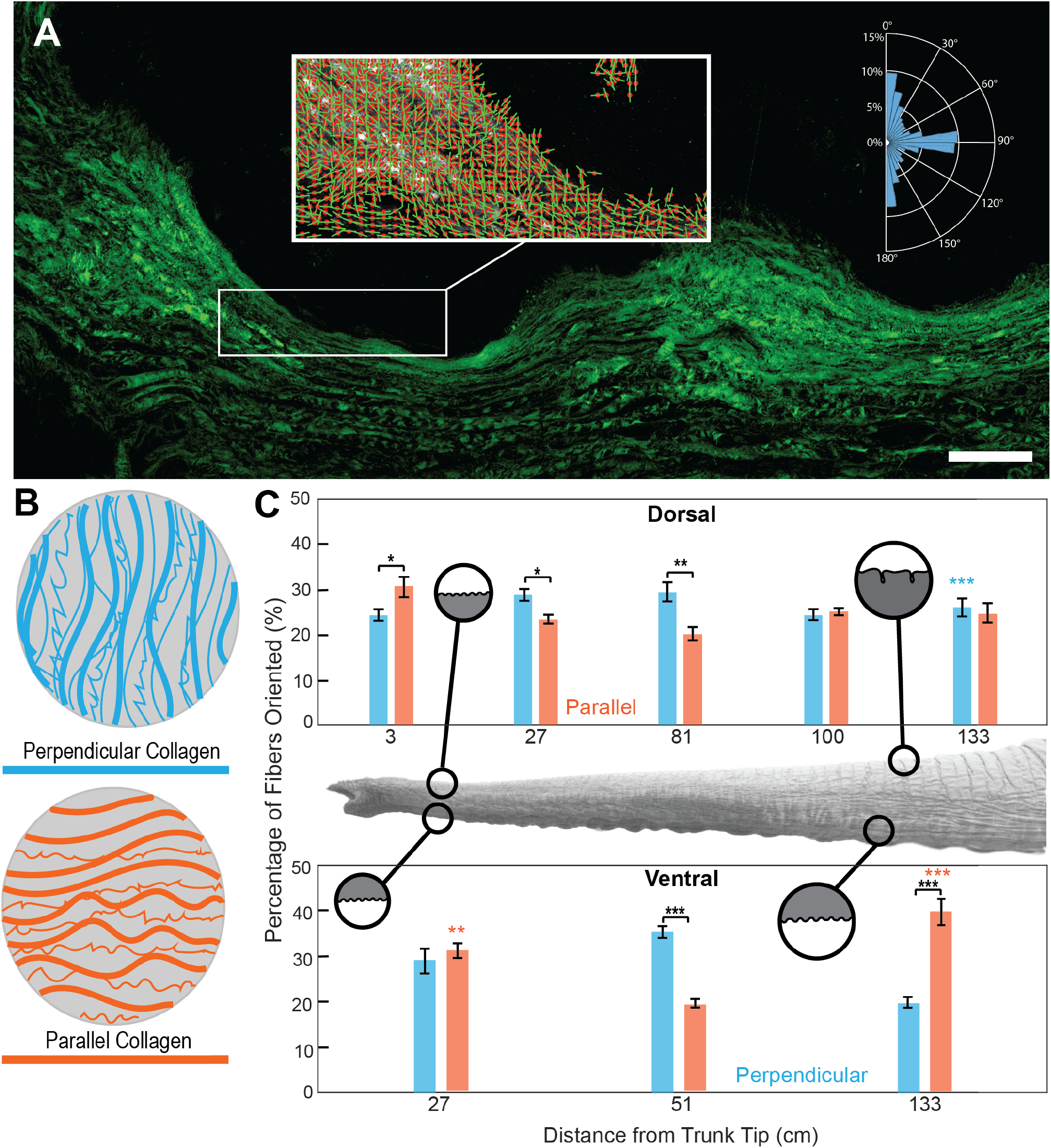
A) Stacked SHG image of the distal ventral elephant trunk with inset of CurveAlign output showing collagen fiber orientation. Inset histogram showing collagen fiber alignment. B) Schematic of parallel and perpendicular collagen fibers in the dermis. C) Relationship between the percentage of collagen fibers and position on the trunk. Parallel fibers are shown in orange and perpendicular fibers in blue. Blue and Orange stars indicate statistical significance between dorsal and ventral sites, with stars placed over the larger value. Black stars indicate statistical significance between perpendicular and parallel comparisons within a single site. Stars indicate the following significance: (^*^*p<* 0.05, ^**^*p <* 0.01, ^***^*p <* 0.001). Scale bar A: 200 *μ*m.

We considered two broad categories of fiber orientation shown in **Figure 5**B. Fibers perpendicular to the skin, shown in blue in the schematic, have angles of 0 *±* 5° and 180 *±* 5°, where 0° is defined as outward normal from the skin as shown in inset of **Figure 5**A. Parallel fibers (orange) have angles of 90*±*5°. To report the number of fibers of these orientations, we report the percentage of fibers oriented in each direction. A histogram of fiber arrangement is constructed and analyzed for the perpendicular and parallel orientation ratios (**Figure 5**A).

We measured the number of collagen fiber overlaps from a dorsal section 133 cm from the tip. The region was a 200 *×* 200 pixel square and an extruded depth of 100 *μ*m. These crossings were counted using ImageJ. In reporting individual collagen fibers, we compared the SHG images of human skin given by Boyle et al. with that of the elephant skin samples in our study (Boyle, Plotczyk, Villalta, Patel, Hettiaratchy, Masouros, Masen, and Higgins, 2019). We measured the average number of overlaps per unit volume and compared this between humans’ plantar and non-plantar tissue and that of elephants.

### Statistical Methods

All calculations, including statistical analysis, were performed with MATLAB 2022A. In the tables and on the figures, values are reported as mean plus or minus standard deviation. We used the MATLAB function *ttest* for t-test to find statistically significant differences between dorsal versus ventral values, difference of values at different positions along the trunk, and differences in perpendicular versus parallel values.

## Results

### Macrostructure of the elephant trunk skin

The outermost layer of the skin, the stratum corneum (SC), is composed of denucleated, keratinized epithelial cells with lipids in between. Underneath the SC is the viable epidermis (VE) which is a sheet of epithelial cells with tight junctions in between them, which gives the skin its barrier function. Beneath this is the structural support layer for the overlying epithelium, known as the dermis (D)(Boyle et al., 2019). To quantify differences in elephant skin morphology across trunk locations, we used H&E image analysis to segment the skin into the SC, the VE, and D (**Figure 2, Figures S1**). Below we will make comparisons of dorsal and ventral skin at the same distance from the tip of the trunk.

Starting with the stratum corneum, we found that on the dorsal trunk, the SC was thickest in the proximal base, with a mean thickness of 2 mm (Table 1, **Figure 3**A), which is significantly different from the ventral SC, with a thickness of 0.34 mm (p*<* 0.001). The remainder of the SC on the dorsal trunk varied from 0.25 mm to 1 mm on average (**Figure 3**A). In contrast, SC of the ventral trunk had a relatively constant thickness of 0.40 mm.

The viable epidermis thickness remained broadly consistent throughout the length of the trunk and between ventral and dorsal sites (Table 1, **Figure S2**). The overall thickness of the VE remained nearly constant at around 0.3-0.4 mm for both the dorsal and ventral elephant surfaces. An exception was the very distal tip of the dorsal skin, 3 cm from the tip (finger at the tip of the trunk), which had a tiny layer of VE at only 0.05 mm thick (**Figure S2**). This thickness displayed a statistically significant difference from the rest of the skin analyzed (p*<* 0.001).

Together, the SC and VE are considered to be the armor for the skin as they serve as the first layers of protection against environmental insults. Compared to other species’ armor layers, such as scales or shells, the elephant skin on the dorsal trunk reaches 2.2 mm thick -this is double the thickness of a pangolin scale and four times that of a human fingernail (**Figure 3**B). Additionally, the epidermal thickness of the elephant trunk is nearly 100 times thicker than the epidermis on an adult human’s torso.

The next skin layer beneath the VE is the dermis. We observe two regions of increased thickness of the dermis, the tip and the proximal base. At the tip, the ventral dermis is 1.5 thicker than the dorsal dermis (2.3 mm versus 1.46 mm thickness, respectively) ((Table 1, **Figure S3**). This thickening makes sense: at the tip, the thicker ventral dermis is where the trunk grasps and manipulates objects. The dermis appears to thicken where the trunk increases in diameter as well. At the proximal base, the dermis along the dorsal trunk is 700% thicker than the dermis at the distal tip (5.44 mm versus 0.8 mm, respectively).

### Micro-structure of elephant skin

To characterize compositional differences in COL1 between the skin samples from the elephant trunk, we used Second harmonic generation imaging (SHG). SHG can identify the macro and micro-level structures of the skin, such as COL1 fiber density, intensity, and orientation **(Figure S4A)**. The color intensity in SHG images can be used as a proxy for fiber strain, indicating the mechanical state of the tissue (Turcotte et al., 2016). (**Figure 4**A-B) showed the ventral trunk has an overall higher intensity than the dorsal trunk, indicating ventral fibers have more pre-strain than dorsal (**Figure 4**C). At the tip of the trunk, the ventral skin has an SHG intensity twice that (*p<* 0.001) seen in the dorsal (**Figure 4**D). This trend was accentuated at the trunk base, where the ventral skin SHG intensity was six times (*p <* 0.001) the intensity of the dorsal skin (**Figure 4**D). The differences in SHG intensity observed here indicate that dorsal skin has less pre-strain imposed on the collagen allowing more stretch-ability than ventral skin.

We next used the SHG images to assess the collagen fiber angle (**Figure S4**, **Figure 5**A). Two fiber angle orientations, perpendicular and parallel relative to the skin surface, are of particular relevance to the physical properties of the skin (**Figure 5**B). As discussed in the methods, we define zero degrees as the outward normal of the skin surface (**Figure 5**A). Perpendicular fibers resist axial trunk loading from forces perpendicular to the skin (**Figure 5**B). The parallel fibers are oriented 90 degrees to the outward normal. Parallel fibers primarily assist with extension and shear loading tolerance (**Figure 5**B).

Upon analysis of the collagen orientation from the SHG images, we found that dorsal skin samples are composed of bi-modal orientation peaks, with COL1 fibers oriented in both the perpendicular and parallel directions (**Figure 5**C). All samples of dorsal skin analyzed have over 20% of perpendicular and 20% of parallel fibers in the skin, indicating a bi-model peak of fiber distribution. Additionally, we see a significant difference when we compare the fiber orientation at specific sites along the trunk. Along the dorsal surface of the trunk at 3, 27, and 81 cm from the tip of the trunk, we see significant differences between the percentage of perpendicular and parallel fibers. The proximal base (100 and 133 cm from the tip) on the dorsal surface, however, shows no significant difference, with around 25% perpendicular and 25% parallel fiber orientation (**Figure 5**C).

Dorsal and ventral surfaces show statistically significant differences in collagen fiber orientation. At the distal tip of the trunk (27 cm from the tip), the ventral skin has significantly more COL1 fibers in the parallel direction (*p <* 0.01) compared to the dorsal skin at the same site (**Figure 5**C). When we look at the proximal base (133 cm from the tip), the dorsal skin has more perpendicular collagen (*p <* 0.001), and less parallel collagen (*p <* 0.001) relative to ventral skin at the same location.

As mentioned above, we see a bi-modal distribution of fiber orientation in the elephant with large percentages in both the perpendicular and parallel directions. In our previous work looking at human skin, we found that both plantar (skin on the sole of the foot) and non-plantar (body) skin contained COL1 fibers with preferential fiber orientation (perpendicular or parallel) in just a single direction(Boyle et al., 2019), as opposed to the bi-model distribution observed in elephant skin. Given the differences in fiber orientation between human and elephant skin, we postulated that there would also be differences in the entanglement of COL1 fibers. To assess COL1 fiber overlap or entanglement (**Figure 6**A), we analyzed a 200 x 200-pixel SHG image segment from dorsal skin 133 cm from the tip. We found that the average number of fiber crossings per *μm*^3^ in the elephant trunk is 5.85 (**Figure 6**B). This value is six times higher than that observed in both human plantar (*p <* 0.01) and non-plantar skin (*p <* 0.01).

**Figure 6:**
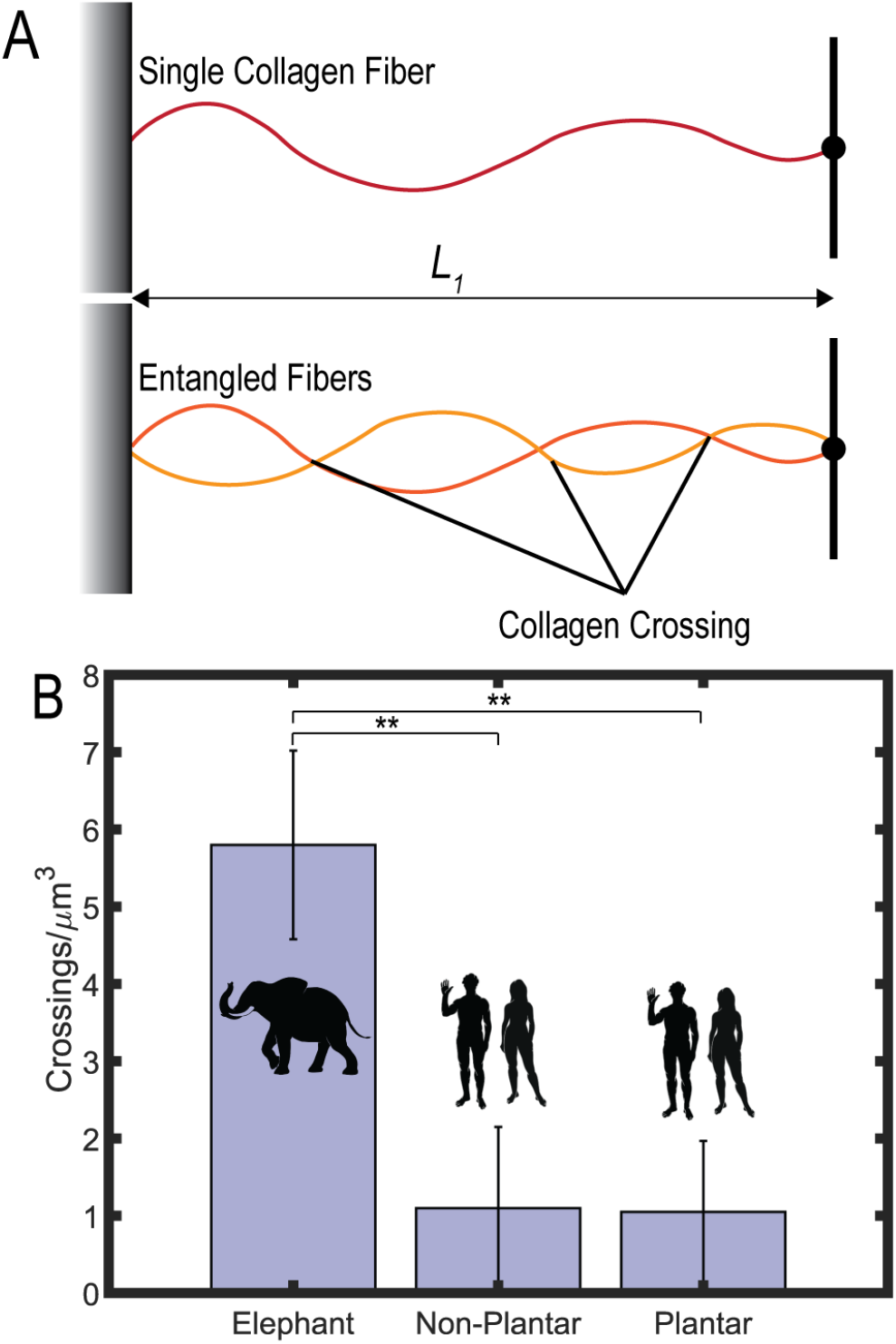
A) Schematic of a cross-linked and non-cross-linked collagen fiber. B) Collagen crossings per cubic micron for elephant skin (dorsal region 133 cm from the tip) and human plantar and non-plantar skin. Published SHG images of human skin reanalyzed from (Boyle et al., 2019). Stars indicate the statistical significance of the difference between elephant and human skin: (^**^ *p <* 0.01) Silhouettes of African elephant (*Loxodonta africana*)from phylopic artist Agnello Picorelli.

## Discussion

We set out to evaluate if elephant trunk skin has variations in its architecture along the length of the trunk that may explain the different functions of the trunk. We found variations in morphology and composition along the trunk length at both the macro and micro scale. The dorsal portion of the trunk, including the trunk’s dorsal finger (3 cm from tip) and dorsal root (133 cm from tip), had the thickest SC layers. The distal tip of the trunk, or finger, is regularly used to manipulate objects, and the dorsal root is more exposed to external stimuli(Dagenais et al., 2021). These functions may explain the thicker dorsal finger and root SC layers.

When we combine the thickness of the SC and VE in this dorsal root and compare it to other species, we see the elephant may have the thickest dermal armor among extant animals; elephants have a dermal armor thickness twice that of a pangolin scale and four times a human thumbnail **Figure 3**B)(Wang, Yang, Sherman, and Meyers, 2016; Wollina, Berger, and Karte, 2001).

While the elephant uses skin for protection, aquatic and arctic species use thick fat layers for protection and insulation (Liwanag, Berta, Costa, Budge, and Williams, 2012). In humans, the sole also has a fat pad that protects the skeleton from heel strike impact. Unlike the fat layers in arctic species, this fat pad does not protect the skin – instead, foot skin has adapted to be thicker and stiffer than body skin, which allows it to withstand mechanical loading. In other species, we see a range of morphological structures, such as shells and scales (**Figure 3**B), where the skin armor has adapted to provide additional protection against environmental pressures (Wang et al., 2016).

Our study was limited by having material samples from only one elephant specimen and one elephant species. While many dry skin samples are available in museums, frozen samples, which allow preservation and histological analysis, are much rarer. Moreover, this specimen was an African bush elephant (*Loxodonta africana*), just one of three elephant species. There may be intrinsic differences between species that we could not address in our study. Asian elephants have only one finger at the tip, with the ventral finger composed of a cartilage bulb. This difference in trunk tip morphology is partly due to Asian elephants being grazers (eat low-lying vegetation). In contrast, African elephants are browsers (also eat high-growing vegetation) and require a prehensile finger to grip and pull leaves off branches for nutrients.

Boyle et al. found that in comparing human skin samples, skin on different body sites had COL1 fibers oriented preferentially in either a parallel or perpendicular direction, depending on the functional requirements for skin at that site (Boyle et al., 2019). The dorsal surface of the elephant trunk expressed relatively even amounts of parallel and perpendicular collagen. The ventral root portion of the trunk had more parallel collagen. We envisage that these observations will give inspiration to future biomimetic studies. While collagen fiber entanglement is still being understood, the general belief is that the structure on the micro-scale leads to unique mechanical responses on the macro scale. There has been increased interest in understanding the macro physical properties that stem from micro-scale entanglements. Such work may influence the design of soft robotic manipulators(Becker, Teeple, Charles, Jung, Baum, Weaver, Mahadevan, and Wood, 2022). Our studies of the impacts of woven fiber structure inside the skin are reminiscent of the impact of patterning in knitted fabric structures. Knitting is a centuries-old activity that involves manipulating a string-like material, traditionally yarn, into a complex fabric with emergent elasticity. These fabrics can exhibit vastly different mechanical properties based on how the stitches, specific slipknots formed by the yarn, are patterned and structured(Singal, Dimitriyev, Gonzalez, Quinn, and Matsumoto, 2023). These structural differences leading to robustness are also challenges in the public health sector. Collagen fibers in skin constructs are always oriented parallel to the skin dermis as they govern how skin contracts. Orienting perpendicular fiber alignment could make skin grafts more robust in their mechanical and flexibility utility.

In summary, we compared the trunk along the distal-proximal and dorsal-ventral anatomical axes, finding differences in the morphology and composition across the elephant trunk and giving insights into the form-function relationships. Elephant trunks have some of the thickest dermal armor in the animal kingdom, with a 2.2 mm thick epidermis. This armor is paired with parallel and perpendicular collagen in the dermis, allowing strength and flexibility. Furthermore, the bi-model orientation of collagen in the dermis leads to individual fiber overlap and interaction, showcasing the entanglement of fibers inside the skin. This work shows the complex nature of elephant skin and provides bio-inspiration for materials that require strength and flexibility.

## Acknowledgements

Thank you to the European Hair Research Society, who supported the travel of AS to Imperial College London. We thank J. Ososky and the Smithsonian Institution Museum of Natural History for assistance with information regarding the frozen elephant trunk. We thank the Facility for Imaging by Light Microscopy (FLIM) at Imperial College London, which is in part, supported by funding from the Wellcome Trust (104931/Z/14/Z) and BBSRC (BB/L015129/1). CH was funded by a project grant from the Engineering and Physical Sciences Research Council (EP/N026845/1).

